# SPNS1 is required for the transport of lysosphingolipids and lysoglycerophospholipids from lysosomes

**DOI:** 10.1101/2022.12.14.520377

**Authors:** Hoa T.T. Ha, Xuan T.A. Nguyen, Linh K. Vo, Nancy C.P. Leong, Siyi Liu, Dat T. Nguyen, Pei Yen Lim, Ya Jun Wu, Toan Q. Nguyen, Jeongah Oh, Markus R. Wenk, Amaury Cazenave-Gassiot, Wei Yi Ong, Long N. Nguyen

**Author notes:** Corresponding author Long N. Nguyen,. **Competing interest**: All authors declare no competing interest. **Classifications**: Biochemistry.

## Abstract

Accumulation of sphingolipids, especially sphingosines, in the lysosomes is attributed to the pathogenesis of several lysosomal storage diseases. In search for a lysosomal protein that mediates the release of sphingosines, we identified SPNS1 which shares the highest homology to SPNS2, a sphingosine-1-phosphate (S1P) transporter. We generated knockout cells and mice for *Spns1* and employed lipidomics and metabolomics to identify SPNS1 ligands. We found that knockouts of *Spns1* resulted in the accumulation of sphingolipids, including sphingosines in embryonic brains and cell lines. These results suggest that deficiency of SPNS1 affects the clearance of sphingolipids in lysosomes. Biochemical assays demonstrated that sphingosines released from lysosomes required SPNS1. Furthermore, by performing a comprehensive analysis of metabolites from livers of postnatal *Spns1* knockout mice (g*Spns1*-cKO), we detected a striking accumulation of lysoglycerophospholipids including LPC, LPE, LPG, and lysoplasmalogens. Interestingly, the release of these lysoglycerophospholipids also required SPNS1. Global knockout of *Spns1* (g*Spns1*-KO) resulted in embryonic lethality between E12.5-E13.5 with developmental defects. Postnatal deletion of *Spns1* in mice caused lipid accumulation in the lysosomes and pathological conditions reminiscent of lysosomal storage diseases. These results reveal a critical molecular role of SPNS1 as a transporter for lysosphingolipids and lysoglyerophospholipids from the lysosomes and link its physiological functions with lysosomal storage diseases.

**Significance:** Phospholipids, including glycerophospholipids and sphingolipids, are delivered to the lysosomes for recycling. The hydrolysis of these lipids by lysosomal enzymes generates the corresponding lysoglycerophospholipids, such as lysophosphatidylcholine and lysosphingolipids, such as sphingosine, which are believed to be exported out of the lysosomes for recycling in the cytoplasm. However, it is unknown how these lysophospholipids are released from the lysosomes. The current study utilized genetic knockout models in combination with mass spectrometry analysis of complex phospholipids and sphingolipids to characterize the roles of an orphan lysosomal transporter, namely SPNS1. These findings show that deficiency of SPNS1 results in the accumulation of lysophospholipids in cells and animal tissues and that the transporter is required to transport both lysoglycerophospholipids and lysosphingolipids out of the lysosomes. SPNS1 is critical for early development in mice. Ablation of SPNS1 at postnatal life causes pathological conditions reminiscent of lysosomal storage diseases in mice. These findings reveal the molecular functions of SPNS1 as a lysophospholipid transporter and provide a foundation for studying the transport of these lysolipids in lysosomal storage diseases.

## Introduction

It is established that lysosomes are the hub for recycling sphingolipids, including sphingomyelins, ceramides, and glycosphingolipids, as well as glycerophospholipids such as phosphatidylcholines and phosphatidylethanolamines. The recycling of sphingolipids appears particularly important for cellular functions because mutation of the enzymes and proteins involved in sphingolipid hydrolysis in the lysosome often results in pathological conditions which are referred to as sphingolipid lysosomal storage diseases (SLSD) (1–3). Several types of SLSD described in the literature, such as Gaucher, Sandhoff, Fabry, Tay-Sachs, and Niemann-Pick type C diseases, remain untreatable (3, 4).

Complex sphingolipids are delivered to the lysosomes via endocytosis or autophagic pathway, to be hydrolyzed by lysosomal enzymes (5). One of the simplest by-products of this process is sphingosine which does not undergo further degradation and must be released from the lysosome for recycling. In the acidic environment of the lysosomes, the amine group of sphingosines is protonated, preventing it from diffusing across the lysosomal membrane into the cytoplasm, where it can be converted into S1P or ceramides. As such, it is hypothesized that sphingosine species are transported out of the lysosomes via a protein-mediated process. However, the identity of the lysosomal transporter for sphingosines is unknown.

Accumulation of sphingosine species has been shown to be detrimental to lysosomal homeostasis and functions (6, 7). In search for a potential lysosomal transporter of sphingosines, we identified Spinster protein homolog 1 (SPNS1), which belongs to a small family of 3 mammalian proteins, namely SPNS1-3. Human SPNS1 shares 57% identities with human SPNS2, which is a plasma membrane transporter for sphingosine-1-phosphate (S1P) (8–10). Additionally, several studies have shown that SPNS1 is expressed in the lysosomes and plays a role in the autophagic pathway (11). Interestingly, SPNS1 was found to be crosslinked with palmitate and sphingosine in cells (12, 13). These reports helped us hypothesize that SPNS1 plays a role in the transport of sphingosines from the lysosome. In this study, we generated the knockouts for *Spns1* in cell lines and mice. We showed that lack of SPNS1 results in the accumulation of sphingolipids such as sphingosines in mouse tissues and cell lines. Additionally, comprehensive metabolomics revealed that lack of SPNS1 also causes the accumulation of lysoglycerophospholipids such as lysophosphatidylcholines (LPC), lysophosphatidylethanolamines (LPE), lysophosphatidylglycerol (LPG), and lysoplasmalogens (lyso-P). We found that SPNS1 is required for the release of sphingosines and lysoglycerophospholipids, such as LPC, from the lysosomes. Importantly, global knockout of *Spns1* in mice resulted in embryonic lethality with severe developmental defects. Furthermore, postnatal knockout of *Spns1* also caused the accumulation of these lysolipids in the lysosomes with pathological conditions resembling SLSD. Our results reveal that SPNS1 plays a critical role in transporting lysolipids out of the lysosomes. The identification of SPNS1 as a lysolipid transporter lays a foundation for future investigations on the transport mechanism of lysolipids from the lysosomes in health and disease.

## Results

### SPNS1 is expressed in the late endosomes and lysosomes and is required for early development in mice

In search for a lysosomal protein with potential transport functions for sphingosines from lysosomes, we identified SPNS1 as a candidate. SPNS1 shares a high sequence identity with SPNS2, a sphingosine-1-phosphate (S1P) transporter from endothelial cells (14), (15), (16). SPNS1 protein is detected in the lysosomal protein fractions (11). To visualize the localization of SPNS1, we developed polyclonal antibodies to detect the expression and localization of SPNS1 in cells. We validated our antibodies using Western blot (**Figure S1A**). By performing immunostaining, we found that SPNS1 is co-localized with LAMP1, a lysosomal marker (**Figure 1A, arrows**). We also detected the expression of SPNS1 in or near the plasma membrane (**Figure 1A, arrowheads**). These data suggest that SPNS1 may be first translocated to the plasma membrane before being destined for the lysosomes. Furthermore, we tested whether SPNS1 is expressed in endocytic vesicles. When SPNS1 was co-expressed with Rab5, an early endosomal marker, we found that they did not co-localized (**Figure S1B**). Interestingly, SPNS1 co-localized with Rab7, a marker for late endosomes (**Figure S1B**). These results suggest that SPNS1 is expressed in late endosomes and lysosomes. In cell lines such as CHO and HEK293 cells, we detected the expression of SPNS1 with a single protein band of approximately 40 kDa (**Figure S1A**). These data suggest that SPNS1 is localized in the lysosomes and that it is also endogenously expressed in cell lines. Lysosome is known for its important roles in the hydrolysis of sphingolipids. The breakdown of these sphingolipids results in sphingosines as one of the final products. The modeled structure of SPNS1 from AlphaFold2 shows that it resembles solute transporters (**Figure S1C**). Therefore, we hypothesize that SPNS1 may be required to release sphingosines from lysosomes. SPNS1 function has not been studied in mouse models. To this end, we deleted *Spns1* in mice and found that a global knockout of *Spns1* (g*Spns1^-/-^*, hereafter g*Spns1*-KO) resulted in embryonic lethality between E12.5-E13.5 with severe defects in the brain and eye (**Figure 1B & S1D**). Viable g*Spns1*-KO embryos were significantly smaller as compared to controls (**Figure 1B-C & S1D**). Nevertheless, the whole anatomy of the gS*pns1*-KO embryos was comparable to that of control embryos (**Figure 1D**). In the brain, we found that the vascularization of the g*Spns1*-KO embryos appeared reduced with increased expression of Glut1 in the neocortices (**Figure 1E-G**). Cortical thickness was significantly reduced (**Figure 1H**). By performing the electron microscopic analysis of the g*Spns1*-KO brain, we found that there was an accumulation of membranous structures in the neural cells (**Figure 1I & S1E**). These results indicate that SPNS1 is required for embryonic development and that its deletion results in the accumulation of membranous structures in the brain.

**Figure 1.**
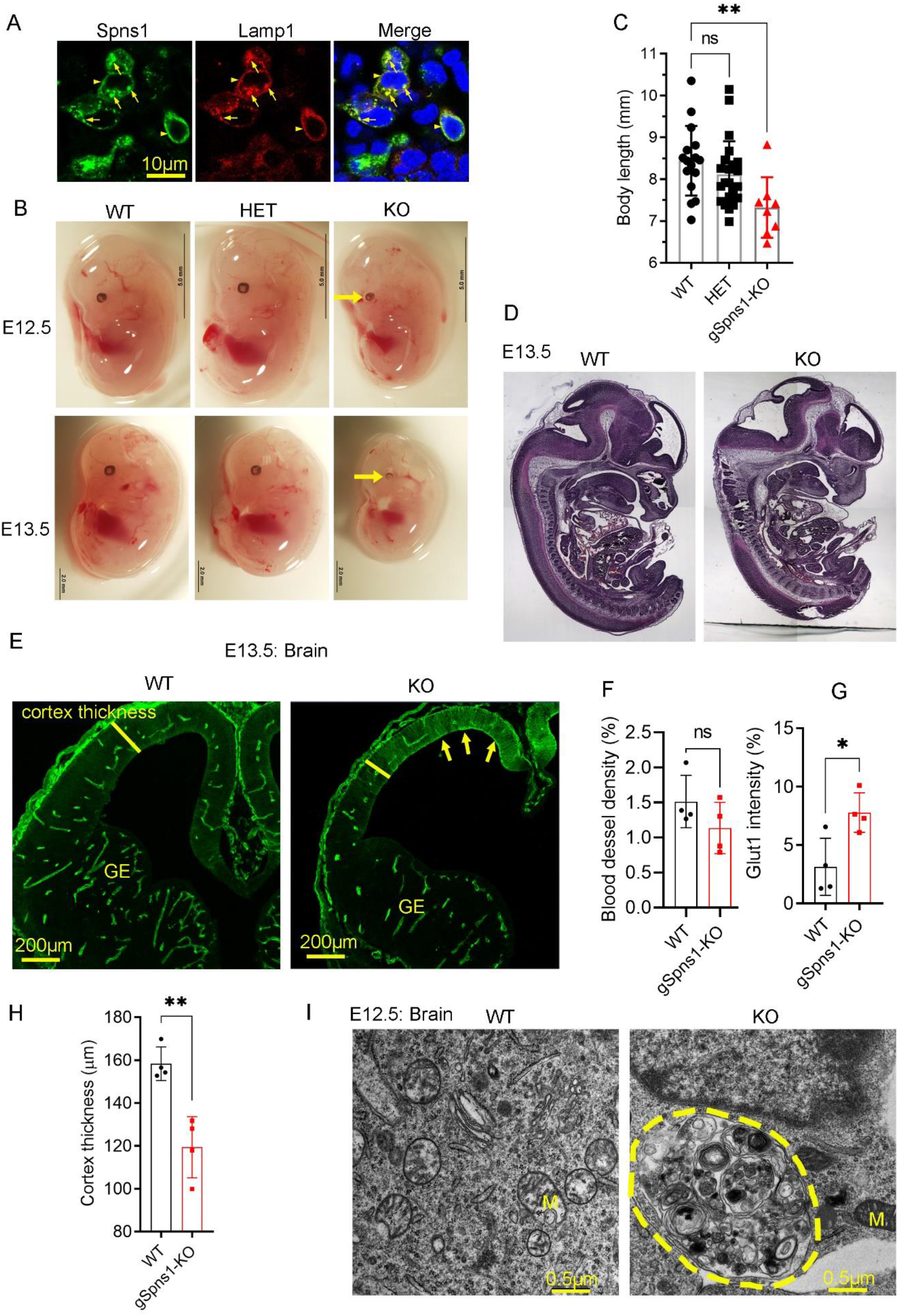
SPNS1 is a putative lysosomal transporter that is required for survival in mice. **A, L**ocalization of SPNS1 in HEK293 cells. Human SPNS1 cDNA was co-transfected with LAMP1-RFP. Spns1 is co-localized with lysosomal marker LAMP1. **B**, Representative images of E12.5 and E13.5 of wild-type (WT), heterozygous (HET), and global Spns1 knockout (gSpns1-KO) embryos. Arrows show maldevelopments of the eyes. **C**, Spns1 KO embryos have smaller body sizes than WT and HET controls. Each symbol represents one embryo. **D**, Gross anatomy of the control and gSpns1-KO embryo at E13.5. **E**, Representative immunostaining of the brain vasculature of controls and gSpns1-KO embryos with Glut1. **F**, Blood vessel density in the brain of control and gSpns1-KO embryo at E13.5. Each symbol represents one embryo. n=4 for each genotype. **G**, Expression of Glut1 in neocortical regions of gSpns1-KO embryos was significantly reduced. Each symbol represents one embryo. n=4 for each genotype. **H**, Cortical thickness of gSpns1-KO embryos was reduced. Each symbol represents one embryo. n=4 for each genotype. **I**, Transmission electron microscopic images of brain sections of E12.5 WT and gSpns1-KO embryos. There were accumulations of membranous structures (demarcated area) in the cytoplasm of brain cells of gSpns1-KO embryos. M, mitochondria. n=3 per genotype. ns, not significant; *p<0.05; **p<0.01. Data are expressed as mean ± SD. Statistical significance was determined by two-sided unpaired t-test.

### SPNS1 is required for sphingosine release from lysosomes

The function of SPNS1 in the lysosome is unknown. The results shown above suggest that lack of SPNS1 causes the accumulation of cellular substances in the brain. To reveal the identities of those substances, we performed lipidomic analysis for sphingolipids of brains from g*Spns1*-KO and control embryos. Interestingly, we found that sphingosine species were significantly elevated in the brains of the g*Spns1*-KO embryos (**Figure 2A; Supplemental table S1**). The levels of ceramides and sphingomyelins were largely unaffected (**Figure 2A**). It is unclear whether sphingosine accumulation was a direct effect of *Spns1* deletion in neural cells of g*Spns1*-KO embryos. To gain further insights, we generated *Spns1* knockouts in HEK293 and CHO cells (*Spns1*-KO cells) (**Figure S2A-B**) and employed the cells for sphingolipid analysis. Consistently, we found that sphingosine species were also accumulated in *Spns1*-KO cells, especially under starvation (**Figure 2B-C; Supplemental table S2-3**). Accumulation of sphingosines has been reported in NPC1-deficient cells. We included NPC1 knockout HEK293 cells (NPC1-KO cells) in our lipidomic analysis for comparison. We found that deletion of *Spns1* resulted in similar accumulation levels of sphingosines compared to NPC1-KO cells (**Figure 2D; Supplemental table S4**).

**Figure 2.**
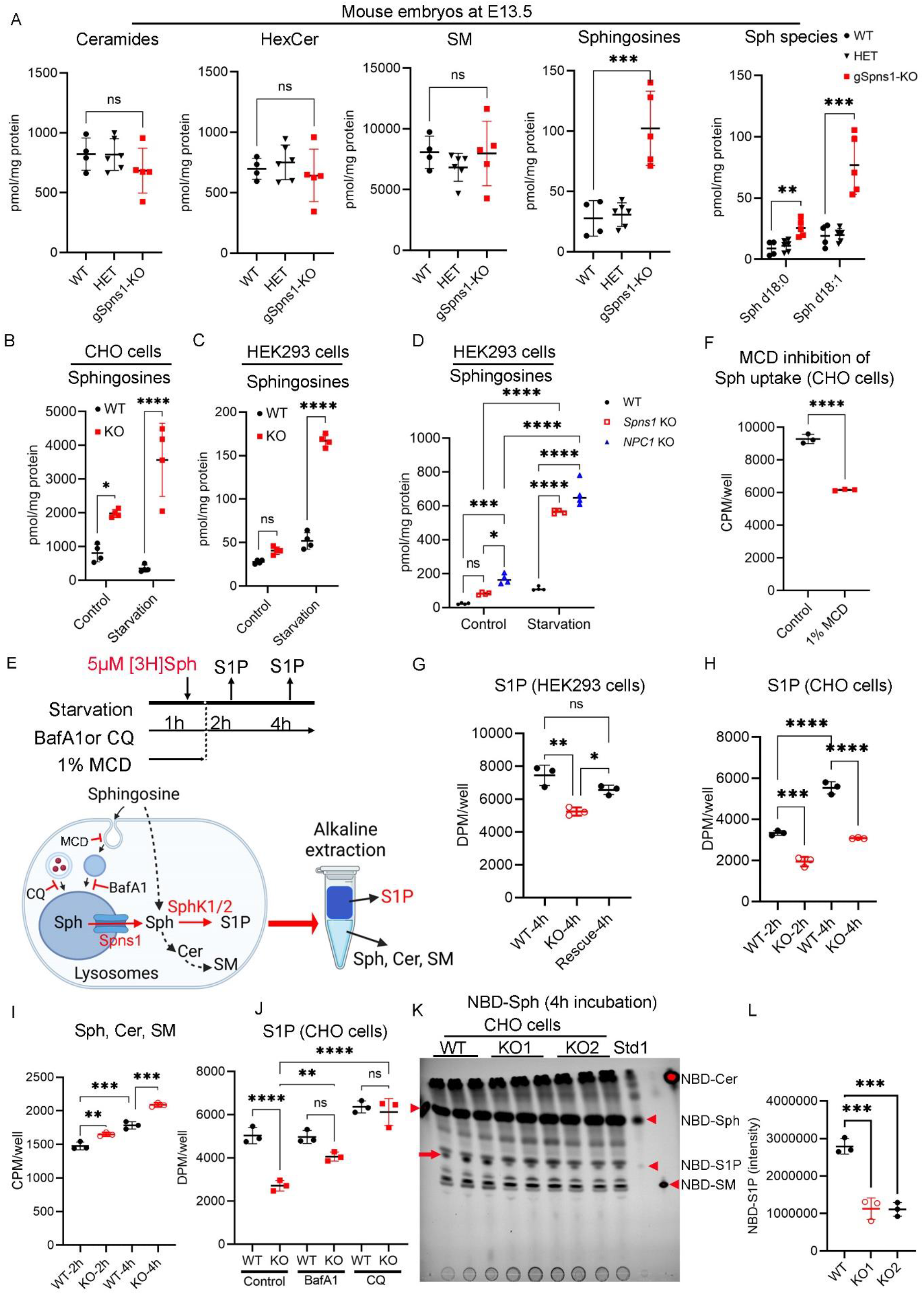
SPNS1 is required for sphingosine release from lysosomes. **A**, Total levels of ceramides, hexosylceramides (HexCer), sphingomyelins (SM), and sphingosines, as well as the levels of individual sphingosine species from whole brains of E13.5 wild-type (WT), heterozygous (HET), and gSpns1-KO embryos. The levels of sphingosine species in the brains of Spns1 knockout embryos were elevated. Each symbol represents one embryo. **B-D**, Total levels of sphingosines from indicated cells with or without starvation condition. Wild-type and Spns1 knockout (KO) CHO cells (in B). Wild-type and Spns1-KO HEK293 cells (in C). Wild-type, Spns1-KO, and NPC1-KO HEK293 cells (in D). Each symbol represents one replicate. **E**, Illustration of [3-^3^H]-sphingosine transport assays. Cells were starved in a medium without amino acids and serum in the presence or absence of bafilomycin A1 (BafA1), chloroquine (CQ), or Methyl-β-cyclodextrin (MCD) for 1 hour. The cells were then added with radioactive sphingosine and incubated until collection for radioactive S1P isolation from other sphingolipids (Sph, Cer, and SM) for quantification. **F**, MCD inhibition of sphingosine uptake in wild-type CHO cells for 1.5 hours. Each symbol represents one replicate. **G**, Radioactive S1P levels (upper phase) from WT, Spns1-KO, and rescue HEK293 cells. Each symbol represents one replicate. **H**, Radioactive S1P levels (upper phase) from WT and Spns1-KO CHO cells. Each symbol represents one replicate. **I**, Radioactive sphingolipid levels (lower phase) from WT and Spns1-KO CHO cells. Each symbol represents one replicate. **J**, Radioactive S1P levels from WT and Spns1-KO CHO cells treated with or without BafA1 and CQ. Each symbol represents one replicate. **K**, TLC analysis of NBD-sphingolipids from wildtype and two different Spns1-KO clones after incubation with NBD-Sph. The arrow shows NBD-S1P band. Arrowheads show standards. Std1 indicates an NBD-Sph/NBD-S1P mixture generated from erythrocytes by incubating with NBD-Sph. **L**, Quantification of NBD-S1P from the TLC plate. *p<0.05; **p<0.01; ***p<0.001; ****p<0.0001; ns, not significant. Data are expressed as mean ± SD. Statistical significance was determined by two-sided unpaired t-test for **F** or one-way ANOVA for **A, B, C, D, G, H, I, J**, and **L**.

The accumulation of sphingosines due to SPNS1 deficiency suggests that SPNS1 plays a role in sphingosine release from the lysosome. Sphingosine can be converted into sphingosine-1-phosphate (S1P) and ceramides in the cytoplasm. Thus, we tested whether the inhibition of SPNS1, which blocks lysosomal sphingosine release, could reduce S1P levels (**Figure 2E**). Exogenous sphingosine is quickly taken up by cells (17). Although it is unclear for the main routes of sphingosine uptake, endocytosis has been shown to deliver sphingosines to the cells (18), (19), (20). Indeed, we validated these findings by incubating the cells with 1% MCD to inhibit endocytosis, significantly reducing sphingosine uptake (**Figure 2F**). Nevertheless, a portion of exogenous sphingosine was also taken up via other routes, presumably by diffusion (**Figure 2E-F**). Taking advantage of the endocytosis of sphingosine, we induced starvation, incubated the cells with [3-^3^H]-sphingosine solubilized in BSA, and examined the levels of [3-^3^H]-S1P synthesized. First, we showed that sphingosine uptake was similar in WT and *Spns1*-KO cells (**Figure S2C-D**). Then, we utilized this assay to screen for different *Spns1*-KO clones from CHO cells. We found that [3-^3^H]-S1P levels in all tested *Spns1*-KO clones were significantly decreased compared to that of WT cells (**Figure S2E-G**). The reduction of S1P levels due to SPNS1 deficiency was also recapitulated in HEK cells, albeit with moderate levels compared to that of CHO cells (**Figure 2G**). Thus, we employed *Spns1*-KO from CHO cells for subsequent experiments. Interestingly, we observed that S1P levels were increased during starvation in WT cells, suggesting that the increased lysosomal activity correlates with the increased release of sphingosine for S1P synthesis (**Figure 2H-I**). Blocking lysosomal activity with bafilomycin A1, an inhibitor of the V-ATPase proton pump, or chloroquine, an inhibitor of lysosome and autophagosome fusion, partially or completely reversed the reduction of S1P levels in *Spns1*-KO cells compared to WT cells (**Figure 2E&J**). Furthermore, using NBD-sphingosine as a substrate, we also observed that NBD-S1P synthesis was significantly reduced in *Spns1*-KO cells compared to that of WT cells (**Figure 2K-L**). These results indicate that SPNS1 is required for sphingosine release from the lysosomes, which is then utilized for S1P synthesis by sphingosine kinases in the cytoplasm.

### Lack of SPNS1 results in the accumulation of lysoglycerophospholipids

We noted that the levels of sphingosine accumulation in *Spns1* knockout embryos were comparable to what has been reported for NPC1 knockouts. However, global knockout of NPC1 is viable, whereas global knockout of *Spns1* is embryonically lethal. Thus, sphingosine accumulation in g*Spns1*-KO embryos is likely the sole cause for the early lethality of *Spns1* knockout embryos. To comprehensively assess the cellular substances accumulated due to the lack of SPNS1, we employed metabolomics to detect metabolites from major metabolic pathways. Thus, we generated conditional knockout mice of *Spns1* using Rosa26Cre-ER^T2^ (hereafter g*Spns1*-cKO mice) (**Figure S3A-B**). Then, we induced *Spns1* deletion at 2 weeks after birth and collected livers for metabolomics (**Figure 3A)**. The metabolomic analysis encompassed over 900 metabolites of all major metabolic pathways, such as amino acids, lipids, nucleotides, carbohydrates, cofactors, and vitamins (**Supplemental table S5-10**). Our data showed that many lipid species were significantly accumulated in the livers of g*Spns1*-cKO mice compared that to controls (**Figure 3B**). The levels of metabolites from the metabolic pathways for amino acids, nucleotides, and carbohydrates, as well as the levels of cofactors and vitamins, were largely unchanged in the livers of g*Spns1*-cKO mice (**Figure S3C; Supplemental table S5-10**). Among the lipid species that were significantly increased, we observed that lysoglycerophospholipids such as LPC, LPE, LPI, LPG, and lysoplasmalogens (including LPC (P) and LPE (P)) were significantly accumulated in the livers of g*Spns1*-cKO mice (**Figure 3C-H**). In contrast, major phospholipids, such as phosphatidylcholine (PC), phosphatidylethanolamine (PE), phosphatidylserine (PS), phosphatidic acid (PA), phosphatidylinositol (PI), and plasmalogens, in the livers of g*Spns1*-cKO mice remained comparable to that of the controls (**Figure S3D; Supplemental table S5-10**). Consistent with the increased levels of sphingosines (**Figure 2 above**), the metabolomic analysis also enabled us to validate the accumulation of most sphingosine species (such as d18:1, d18:0, d16:1 sphingosine species) in g*Spns1*-cKO mice (**Figure 3I & J**). Ceramides and lactosylceramides but not sphingomyelins were also significantly accumulated in the livers of g*Spns1*-cKO mice (**Figure 3K; S3D**). The levels of cholesterol and its related metabolites in the livers of the g*Spns1*-cKO mice was not elevated (**Figure S3D; Supplemental table S5-10**). We validated the accumulation of these lysoglycerophospholipids and lysosphingolipids by targeted lipidomics in additional animals (**Figure S4**). Therefore, these data indicate that postnatal deletion of *Spns1* in mice not only results in the accumulation of lysosphingolipids but also lysoglycerophospholipids. To gain further insights into the potential role of SPNS1 as a transporter for the lysoglycerophospholipids, we incubated *Spns1*-KO and wild-type CHO cells with NBD-palmitate and analyzed NBD-labelled phospholipids by thin layer chromatography (TLC) (**Figure 3L-N**). Notably, we found that NBD-LPC was significantly accumulated in *Spns1*-KO cells compared to control cells (**Figure 3M-N**). These findings indicate that deletion of *Spns1* also results in the accumulation of lysoglycerophospholipids such as LPC.

**Figure 3.**
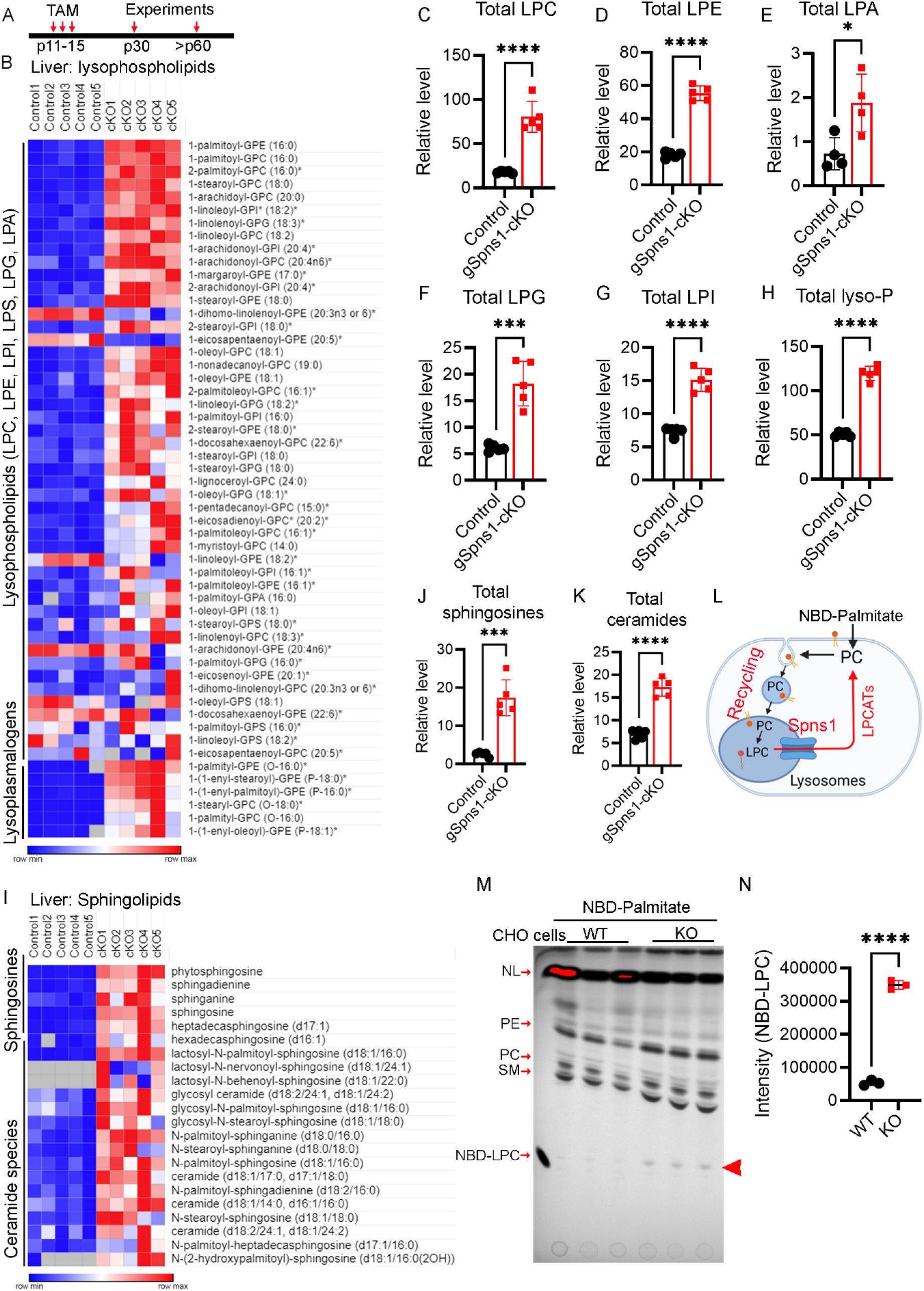
Comprehensive metabolomic analysis reveals the accumulation of lysoglycerophospholipids and lysosphingolipids in the livers of gSpns1-cKO mice. **A**, Illustration for postnatal deletion strategy of Spns1 using Rosa26Cre-ER^T2^ mice. **B**, Heatmap of lysoglycerophospholipids from livers of controls and gSpns1-cKO mice. n=5 mice per genotype. **C-H**, Total levels of lysophosphatidylcholine (LPC), lysophosphatidylethanolamine (LPE), lysophosphatidic acid (LPA), lysophosphatidylglycerol (LPG), lysophosphatidylinositol (LPI), lysoplasmalogens (lyso-P). Each symbol represents one mouse. **I**, Heatmap of sphingolipids from livers of controls and gSpns1-cKO mice. n=5 mice per genotype. **J**, Total levels of sphingosines from livers of control and gSpns1-cKO mice. n=5 mice per genotype. **K**, Total levels of ceramides from livers of control and gSpns1-cKO mice. n=5 mice per genotype. **L**, Illustration of NBD-palmitate labeling experiment. **M**, Thin layer chromatography analysis of NBD-labelled phospholipids after 24 hours of pulse-labeling with NBD-palmitate in WT and Spns1-KO CHO cells. Arrowhead indicates NBD-LPC band. NL, neutral lipids. **N**, Quantification of NBD-LPC from the TLC plate. *p<0.05; ***p<0.001; ****p<0.0001; ns, not significant. Data are expressed as mean ± SD. Statistical significance was determined by two-sided unpaired t-test.

### Deficiency of SPNS1 results in phenotypes reminiscent of lysosomal storage diseases in mice

Global knockout of *Spns1* results in early embryonic lethality, indicating that SPNS1 plays a critical role in the development. To gain further insights into the physiological roles of SPNS1, we characterized the conditional knockout mice of *Spns1* after birth (**Figure 4A-F**). Postnatal deletion of *Spns1* results in a significant reduction of body weight in both male and female mice 2 weeks post-injection of tamoxifen (**Figure 4B-C**). Increased white blood cell counts were reported in models of lysosomal storage diseases (21), (22). We found that white blood cell (WBC) count of g*Spns1*-cKO was significantly elevated (**Figure 4D**). Interestingly, we observed that the livers of g*Spns1*-cKO mice exhibited hyperplasia (**Figure 4E-F**). Histological assessment of liver sections revealed that hepatocytes exhibited foamy phenotype, a sign of lipid accumulation that has been observed in lysosomal storage diseases (**Figure 4G**). We found that expression of the Cathepsin B was clustered around the nucleus with increased fluorescent intensity in hepatic cells of g*Spns1*-cKO mice (**Figure 4H-I, arrows**). Furthermore, we found that there were increased numbers of macrophages in the livers of g*Spns1*-cKO mice compared to controls (**Figure 4J**). Interestingly, we found that the maturation of the Cathepsin B protein was defective (**Figure 4K**), suggestive of lysosomal defects (23). In addition, expression of LC3B-II was slightly increased in livers of g*Spns1*-cKO mice compared to controls (**Figure 4L&M**). These results point to a possibility that pathological conditions in the livers of g*Spns1*-cKO mice could be linked to a defect in lipid clearance from the lysosomes. Thus, we enriched lysosomes from the livers of g*Spns1*-cKO and control mice and performed lipidomic analysis for lysosphingolipids and lysoglycerophospholipids (**Figure S5A-B**). We found that these lysophospholipids were significantly accumulated with high levels in lysosomal fractions from livers of g*Spns1*-cKO mice (**Figure 4N-O, Figure S5C, Supplemental table S11**). These results reaffirm that SPNS1 is indispensable for the transport of these lysophospholipids out of the lysosomes. Collectively, our findings indicate a deficiency of SPNS1 results in lipid accumulation in the lysosomes and pathological conditions that resemble lysosomal storage diseases.

**Figure 4.**
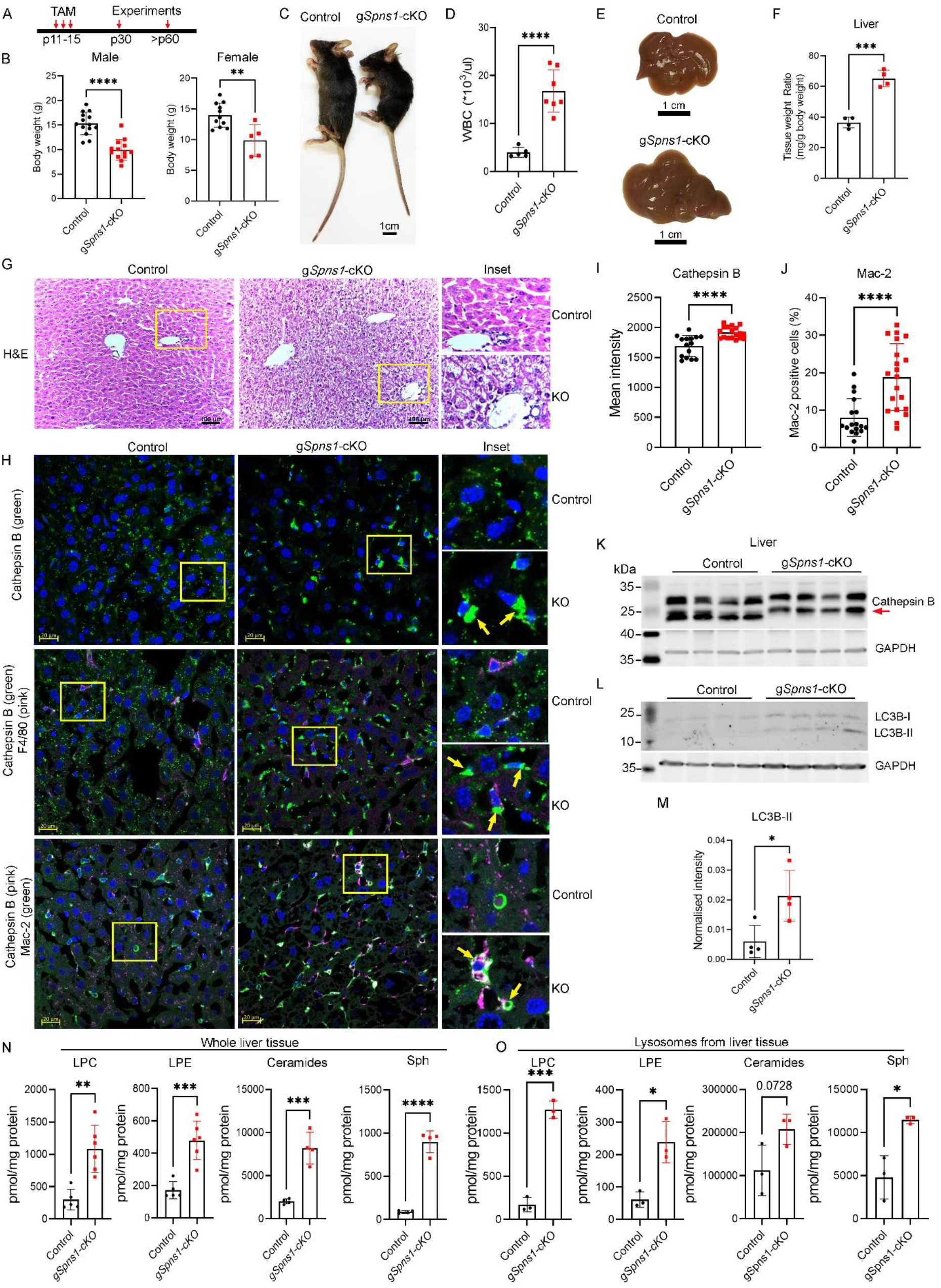
Spns1 knockout mice exhibit lysosomal storage phenotypes. **A**, Illustration for postnatal deletion strategy of Spns1 using Rosa26Cre-ER^T2^ mice. **B**, Reduction of body weights in gSpns1-cKO female and male mice after 2 weeks of tamoxifen treatment. Each symbol represents one mouse. **C**, Representative images of a control and a gSpns1-cKO mouse. **D**, Increased number of white blood cells (WBC) in gSpns1-cKO mice compared to controls. Each symbol represents one mouse. **E**, Representative images of livers from a control and a gSpns1-cKO mouse. **F**, Increased liver weights in gSpns1-cKO mice compared to controls of the same ages. Each symbol represents one mouse. **G**, Representative images of H&E staining of liver sections from control and gSpns1-cKO mice. Inset image shows the foamy phenotype in the livers of gSpns1-cKO mice. n=3 per genotype. **H**, Representative images of immunostaining of liver sections from control and gSpns1-cKO mice with Cathepsin B or with co-staining of Cathepsin B and F4/80 for Kupffer cells or with Cathepsin B and Mac-2 for macrophages. n=3 per genotype. **I**, Quantification of Cathepsin B fluorescent intensity from H**. J**, Quantification of the number of Mac-2 positive cells from H. Each symbol represents one section from n= 3 mice per genotype. **K-L**, Western blot analysis of Cathepsin B and LC3B from whole liver protein lysates of control and gSpns1-cKO mice. Cathepsin B processing was defective in the livers of gSpns1-cKO mice (arrow). n=4 per genotype. **M**, Quantification of total LC3B-II bands from L. **N**, Total levels of LPC, LPE, ceramides, and sphingosines from whole livers of control and gSpns1-cKO mice. n=4-6 per genotype. **O**, Total levels of LPC, LPE, ceramides, and sphingosines from lysosomal fractions (F1 and F2) of the livers of control and gSpns1-cKO mice. n= 3 per genotype. *p<0.05; **p<0.01; ***p<0.001; ****p<0.0001; ns, not significant. Data are expressed as mean ± SD. Statistical significance was determined by two-sided unpaired t-test.

## Discussion

SPNS1 was first discovered in flies as a gene involved in mating behaviors (24). The mutant flies of the same gene (namely *Spin*) exhibited aberrant apoptosis of neurons and glial cells (24). Interestingly, SPIN mutant flies show an accumulation of auto-fluorescent, lipofuscin-like pigments in the brain at the early stage of development (22, 24–26). Lipidomic analysis of the SPIN mutants showed that ceramides and sphingosines were significantly accumulated in the brain (25). In zebrafish, deletion of *Spns1* resulted in lethality. SPNS1 mutant fishes exhibited an accumulation of lipofuscin-like structures in the muscle and liver and accelerated signs of aging and shorter lifespan (26). The mutant fishes also exhibited delayed clearance of autophagosomal contents (27). Consistent with the essential role of SPNS1 for survival from flies to fishes, we showed that global knockout of *Spns1* in mice results in embryonic lethality between E12.5 and E13.5.

The molecular function of SPNS1 is rather elusive, and the nature of the lipofuscin phenotypes in animals with SPNS1 deficiency was uncharacterized. Several other studies have suggested that SPNS1 is involved in transporting sugars and amino acids (28, 29). However, these studies fail to explain the nature of lipofuscin structures and lipid accumulation in SPNS1-deficient cells and animals (25). We found that knockout of *Spns1* in mice results in the accumulation of sphingolipids, especially sphingosines. Sphingosine is protonated in the lysosomal lumen’s acidic compartment, preventing it from diffusing via the membranes. Mechanistically, our results point out that SPNS1 is required for the release of lysosomal sphingosines for recycling in the cytoplasm. Recently, SPNS1 was reported as a transporter for LPC and LPE from the lysosomes (30). Indeed, we also show that the lack of SPNS1 results in remarkably elevated levels of lysoglycerophospholipids such as LPC, LPE, LPG, and lysoplasmalogens. Our biochemical experiments also demonstrate that the lack of SPNS1 causes the accumulation of LPC in the cells. Thus, these findings indicate that SPNS1 is also responsible for the release of lysoglycerophospholipids from the lysosomes. Interestingly, both our data and the data from He et al. show that lack of SPNS1 causes elevated levels of sphingosines and ceramides in mice (30). While we were able to show that SPNS1 is required for sphingosine transport from the lysosomes for recycling, He et al., based on the cell surface transport assays, ruled out this molecular function of SPNS1. Exogenous sphingosine is rapidly taken up via endocytosis (18–21, 31). We argue that the authors of the study underestimated the role of endocytosis in their cell surface transport assays for sphingosine.

Thus, ruling out the sphingosine transport function of SPNS1 by the study was unproven. Although the direct role of sphingosine transport remains to be established, our results are supported by the previous reports that cross-linkable sphingosine binds to SPNS1 (12, 13).

At this point, the lack of SPNS1 results in the accumulation of both lysoglycerophospholipids and lysosphingolipids in the lysosomes. However, it is unclear what is the major cause that leads to early lethality and phenotypic changes observed in conditional knockout mice of *Spns1*. Complex glycerophospholipids such as PC and PE are hydrolyzed by lysosomal phospholipase A2, mainly PLA2G15 (Also known as LPLA2), to liberate respective lysoglycerophospholipids (32). However, lack of the lysosomal hydrolysis of these glycerophospholipids appears less detrimental to the mice. Indeed, mice lacking PLA2G15 that results in the accumulation of phospholipids, but not sphingolipids in the lysosomes are viable and only exhibit phenotypes at later ages (32, 33). It is also worth noting that lysoglycerophospholipids can be further hydrolyzed into free fatty acids and glycerophosphodiesters such as GPC and GPE in the lysosomes. Indeed, a recent study showed that CLN3, a gene mutated in Batten disease, is shown to be a transporter for these glycerophosphodiesters from lysosomes, indicative of an active transport pathway for these compounds (34). Our metabolomic results show that lack of SPNS1 does not cause accumulation of these glycerophosphodiesters, implying that the release of these by-products from hydrolysis of lysoglycerophospholipids is still active in *Spns1* knockout cells. Thus, it seems that the accumulation of lysoglycerophospholipids in the lysosomes causes milder pathological conditions compared to that of sphingolipids. We believe that the accumulation of sphingolipids, especially sphingosines, due to lack of SPNS1, might be a major causal factor. Indeed, these sphingolipids have been shown to be detrimental to the cells and animal models, notably in models of sphingolipid lysosomal storage diseases (7), (2), (35). In line with the essential role of sphingosine export from the lysosome, knockout of acid ceramidase (Asah1), the lysosomal ceramidase in mice causes embryonic lethality (36). Deficiency of Asah1 in adult mice also results in severe phenotypes reminiscent of phenotypes of conditional knockout of *Spns1* mice (22). These findings imply that the release of sphingosines from the hydrolysis of complex sphingolipids in the lysosomes is a critical step to avoid lysosomal defects. Nevertheless, it remains to be determined for the toxicity caused by the accumulation of each type of the lysolipids in the lysosomes of SPNS1-deficient cells.

In summary, our findings show that SPNS1 is required for the transport of lysosphingolipids as well as lysoglycerophospholipids from the lysosomes. Furthermore, the identification of SPNS1 as a transporter for these lysolipids provides a foundation for future study on the transport mechanisms of these lipids from the lysosomes.

## Material and methods

### Mice

All the mice were in C57BL/6 background. The whole-body knockout of *Spns1* was obtained from International Mouse Phenotyping Consortium (IMPC). To generate the global knockout embryos (*Spns1^−/-^*; hereafter g*Spns1*-KO), heterozygous (*Spns1^+/−^*) *Spns1* mice were inter-crossed, and the embryos were collected from timed-mated pregnant females. On the day of the experiments, pregnant mice were anesthetized by CO2, and then the embryos were collected. The wild-type, heterozygous, and homozygous knockout embryos of the same litter were collected for experiments. Tail tips were collected for genotyping. The genotyping primers are listed in the supplementary.

In order to generate the conditional knockout of *Spns1* (*Spns1^f/f^*), two loxP sites were inserted on both sides of the exon 2 using CRISPR/Cas9 technology. The mice were generated by the Transgenic and Gene Targeting Facility (TGTF), at the Cancer Science Institute of Singapore (CSI). To generate the postnatal deletion of *Spns1* in the whole-body (hereafter g*Spns1*-cKO), conditional *Spns1^f/f^* mice were bred with Rosa26Cre-ER^T2^ mice. The Cre expression was induced by intraperitoneal injection (IP) of tamoxifen (TAM) at a dose of 50 mg/kg body weight at P11, P13, and P15. Tamoxifen was prepared by dissolving in corn oil with a 20 mg/ml concentration. All animal procedures were reviewed and approved by IACUC protocols (BR19-0633 and R019-567).

### Cell culture

HEK293 cells and CHO cells were cultured in DMEM (Gibco, Cat 11995065) media supplemented with 10% fetal bovine serum (Gibco, Cat 10270106) and 1% Penicillin/Streptomycin (Gibco, Cat 15140122) in the 37 °C incubator with 5% CO2. The cells were starved in the DMEM medium without FBS and without amino acid (Wako, 048-33575).

### CRISPR/Cas9 gene deletion

*Spns1* coding sequence in HEK293 and CHO cells was targeted by CRISPR/Cas9 technology. Briefly, individual gRNA generated using CRISPR/Cas9 v2 plasmid (Addgene: 52961) was transfected to HEK293 or CHO cells. The following gRNAs were used: 5’-ATTCCTGCTGCGTTCCCGCGTGG-3’ (for HEK293 cells), 5’-GCACGAGATCCCGGACCGCGAGG-3’ and 5’-TGATGCGCTGCAGACCCTCGCGG-3’ (for CHO cells). After 24 hours of transfection using lipofectamine 2000 (Invitrogen), single cells were sorted by flow cytometry onto 96-well plates. After expansion, these *Spns1* knockout (KO) clones were subjected to Western blot analysis to detect SPNS1 protein using in-house generated polyclonal antibodies. Clones with a complete absence of SPNS1 protein band were selected for experiments. To rescue SPNS1 in *Spns1*-KO HEK293 cells, full-length *Spns1* cDNA was knocked in *Spns1* locus using CRISPR/Cas9 technology. Expression of SPNS1 was confirmed by Western blot in rescue cells.

### Enrichment of lysosomes from liver tissues

Lysosomes from mouse livers were isolated using the tyloxapol method that has been previously described (37). The detail protocol is provided in the supplementary. Briefly, control and g*Spns1*-cKO mice were injected with 4 μl/g body weight of a 17% (w/v in 0.9% NaCl) Triton WR1339 solution (Sigma Aldrich) for three days prior to fractionation of lysosomes. The non-perfused livers were homogenized in 0.25 M sucrose, and then centrifuged to collect the post-nuclear supernatant (PNS). Lysosomes were extracted from PNS by sucrose gradient using ultracentrifuge. The first two fractions containing lysosomes were collected to analyze by western blot and lipidomic.

### Immunostaining and confocal microscopic analysis

Embryos were collected at E12.5 and E13.5 from timed-mated mice and then fixed in 4% PFA prepared in PBS at 4°C overnight. The next day, embryos were changed to 15% sucrose for 24 hours and then 30% sucrose for 24 hours prior to being embedded in OCT. Embryonic brains were dissected into 25-30 μm sections for immunostaining (IF). Briefly, sections were washed twice with PBS for 30 minutes. Then, sections were permeabilized in 0.1% Triton X-100 in PBS for 2 hours, followed by the incubation with a blocking solution containing 1% BSA, 0.3% Normal goat serum (NGS), and 0.1% Triton X-100 in PBS for 1 hour. Next, sections were incubated with anti-mouse GLUT1 (Abcam, ab40084,1:200) at 4°C overnight, followed by 3 washes in PBS at 5-minute intervals. Then, the goat anti-mouse antibody (Alexa Fluor 488, Thermofisher A11034,1:500) was added to the slides for 1-hour incubation at room temperature to visualize the GLUT1 signal. The sections were washed twice with PBS (5-minutes interval) and then stained with Hoechst 33342 (Thermofisher, #62249, 1:1000 in PBS) for 15 minutes at room temperature. After washing the slides twice with PBS, the sections were mounted with mounting media. The sections were imaged using Zeiss LSM710 confocal microscope. Similarly, immunostaining for liver sections from control and g*Spns1*-cKO mice was also performed (details provided in supplemental information). For visualization of lysosomes, Cathepsin B antibody was used. For visualization of Kupffer cells and infiltrated macrophages, F4/80 and Mac-2 were used, respectively. The immunofluorescence quantification method is described in the supplementary methods.

### Sphingosine transport assay with and without inhibition of autophagy/endocytosis

For sphingosine transport assays, wild-type HEK293, CHO, or corresponding Spns1-KO cells were used. Before transport assays, the confluent cells were starved in DMEM medium without amino acids and serum (Wako, 048-33575) for 1 hour. Next, the cells were incubated with 5 μM [3-^3^H]-Sph (American Radiochemicals) prepared in 12% BSA in the same starvation medium for 2 or 4 hours. The cells were washed once with DMEM medium containing 0.5% BSA and lyzed in RIPA buffer for lipid extraction. [3-^3^H]-S1P was then isolated from the cell lysates using alkaline extraction methods (38, 39). This method recovers approximately 95% S1P in the upper phase. The radioactive signals in the upper phase (containing [3-^3^H]-S1P) and lower phase (containing [3-^3^H]-sphingosine, [3-^3^H]-ceramides, [3-^3^H]-SM) were quantified by a scintillation counter (Perkin Elmer Tri-Carb Liquid Scintillation Analyzer).

For sphingosine transport assays with Methyl-β-cyclodextrin (MCD) inhibition, WT CHO cells were incubated in starvation medium containing 1% MCD (w/v), for 1 hour. The cells were then incubated with 5 μM [3-^3^H]-Sph prepared in 12% BSA in the same starvation medium containing 1% MCD for 30 minutes. The cells were washed once with plain DMEM medium containing 0.5% BSA and lyzed in 500 μl RIPA buffer for scintillation quantification.

For bafilomycin A1 and for chloroquine inhibition, the confluent cells were incubated in starvation medium containing 500 nM bafilomycin A, or 0.1 mM chloroquine for 1 hour. Next, the cells were incubated with 5 μM [3-^3^H]-Sph (American Radiochemicals) prepared in 12% BSA in the same starvation medium with and without the drugs for 4 hours. Isolation of [3-^3^H]-S1P was done prior to the quantification by a scintillation counter.

### NBD-Sphingosine and NBD-Palmitic acid labeling assay

For NBD-sphingosine labeling, wild-type and *Spns1*-KO CHO cells were incubated with 10 μM NBD-sphingosine (Cayman, 1449370-25-5) dissolved in 12% BSA for 4 hours under starvation condition. Cells were washed once with PBS. For NBD-palmitate labeling, 10 μM NBD-palmitate (Avanti, 810105P) solubilized in 12% BSA was added to the cells for 24 hours in the growth medium. Lipids from cells were extracted twice with HIP buffer (Hexane : 2-propanol, 3:2, v/v), as described in the thin layer chromatography (TLC) method, prior to separation in a TLC plate.

### Thin Layer Chromatography (TLC)

Lipids from cells were extracted twice with 500 μl HIP buffer and dried under nitrogen gas. Sphingolipids were separated in TLC by solvent 1-butanol/acid acetic/water (3:1:1, v/v). Phospholipids were separated in a TLC plate using chloroform/methanol/H2O (65:25:4, v/v). Fluorescent lipids on TLC plates were visualized by ChemiDoc (Bio-rad) and quantified using ImageLab.

### Hematological analysis

Blood samples were collected into EDTA-K2-coated tubes from control and g*Spns1*-cKO mice and directly analyzed using Celltex-a MEK-6400 (Nihon Kohden).

### Metabolomics

For metabolomics, we collected 50-100 mg tissues from PBS-perfused livers of *Spns1^f/f^*-Rosa26Cre-ER^T2^ (g*Spns1*-cKO) and control mice. The untargeted metabolomics was performed by Metabolon. Detail methods for metabolomic analysis from the company’s description are described in the supplemental information. Heat maps of differential levels of metabolites were generated using Morpheus, https://software.broadinstitute.org/morpheus.

### Transmission electron microscope

Wild-type (n=3) and global knockout embryos of *Spns1* (n=3) at E12.5-E13.5 were collected and fixed in 2.5% glutaraldehyde for 24 hours at room temperature. Mouse embryo sections were prepared and were post-fixed with 1% osmium tetroxide for 1 hour and dehydrated using a serial ethanol concentration. Images were taken using a Fei Quanta650 transmission electron microscope.

### Statistics analysis

Data were analyzed using GraphPrism9.0 software. Statistical significance was calculated using the student’s t-test (two-tailed), Welch’s t-test, and one-way or two-way ANOVA for multiple comparisons. P-value < 0.05 was considered as significant and indicated by asterisks. P-value > 0.05 was indicated as not significant (ns).

## Acknowledgement

We thank Ms Brenda Lam for helping with H&E staining. We are grateful to Dr. Roberto Zoncu from UC Berkeley for the gift of NPC1 knockout cells. This study was supported in part by Singapore Ministry of Health’s National Research Council NMRC/OFIRG/0066/20, Ministry of Education T2EP30221-0012, NUHSRO/2022/067/T1, and NUSMED-FOS Joint Research Programme on Healthy Brain Aging grants (to L.N.N.).

