## Supplementary information for "SPNS1 is required for the transport of lysosphingolipids and lysoglycerophospholipids from lysosomes"

**This file contains:**

- Supplementary methods
- Figure and figure legend S1-S5
- Legends for supplemental table S1-S14

Other supplementary data for this work include:

- Datasets S1-S14

**Supplementary methods**

**Antibodies.** Primary antibodies including LAMP1 (sc-20011) and GAPDH (sc-32233) were from Santa Cruz Biotechnology. Calreticulin (CST-12238), LC3B (3686S), and Cathepsin B (31718S) antibodies were from Cell Signaling Technologies. GLUT1 (ab40084) antibody was from Abcam. Mac-2 (125401) and F4/80 biotin-conjugated (123105) antibodies were from Biolegend. Secondary antibodies including Donkey anti-rabbit 800 (926-32213), Donkey anti-rabbit 680 (926-68023), and Goat anti-mouse 680 (926-68020) antibodies were from Licor company. Goat anti-mouse (Alexa Fluor 488, A11034) was from Thermofisher. Goat anti-rabbit (Alexa Fluor 488 (A11034), Alexa Fluor 633 (A21071)), and goat anti-rat (Alexa Fluor 488 (A11006)) antibodies were from Invitrogen. Streptavidin (Alexa Fluor 647, 016-600-084) antibody was from Jackson ImmunoResearch. Antibodies for human and mouse SPNS1 were generated in-house using C-terminal epitope DDRIVVPQRGRSTRVPV.

**Sphingosine uptake assay**. For whole-cell sphingosine (Sph) uptake assays, the WT and Spns1-KO CHO cells were incubated in the transport buffer containing 5 µM [3-^3^H]-Sph for 15 minutes. Transport buffer contains 0.54 mM KCl, 1.3 mM CaCl2, 0.53 mM MgCl2, 0.4 mM MgSO4, 0.37 mM KH2PO4, 138 mM NaCl, 0.28 mM Na2HPO4, 5.5 mM glycine, 5.6 mM D-glucose adjusted with HCl to pH 5. The cells were washed once with plain DMEM medium containing 0.5% BSA and lyzed in 500µl RIPA buffer for scintillation quantification.

**Genotyping.** The following primers were used for genotyping the global Spns1 knockout: 5’-GGTAGAGCCAGGTGTGTTGGC-3’ (forward primer), 5’-GATCTAACCCACCTCCTTCCTTTCC-3’ (reverse primer 1), 5’- GGGCAAGAACATAAAGTGACCCTCC-3’ (reverse primer 2). For genotyping Spns1f/f mice, the following primers were used: 5’-GTAGAGTGGGCAGGGTAATGTG-3’ (forward primer F1), 5’-GGATGGTGCGACATCAGTGA-3’ (reverse primer R1), that generated an WT band with 448 bp and a floxed band with 482 bp. Deletion of Spns1 produced a 135 bp band. For genotyping the Cre-ERT2, these primers were used: 5’-AAGGGAGCTGCAGTGGAGTA-3’ (common forward primer), 5’-CGGTTATTCAACTTGCACCA-3’ (mutant reverse primer), 5’-CCGAAAATCTGTGGGAAGTC-3' (wild-type reverse primer), that generated a Cre band of 450 bp (mutant) and an WT band of 297 bp.

**Immunostaining of mouse liver sections.** For immunostaining of the liver sections from adult mice, 10 µm frozen sections were washed in warm PBS for 30 minutes twice to remove OCT completely. Permeabilization was then performed in 0.5% Triton X-100 in PBS (PBST) for 1h followed by blocking in 5% NGS in PBST. Sections were incubated with anti-Cathepsin B (Cell Signaling, 31718S, 1:200) or co-stained with anti-Mac 2 (Biolegend, 125401, 1:250) or anti-F4/80 (biotin-conjugated, Biolegend, 123105, 1:500) in blocking buffer at 4°C overnight. All sections were washed thrice in PBS at 5 minutes interval. Depending on different combinations, the sections were incubated with dilution 1:500 of secondary antibodies goat-anti rabbit (A488 or 633), goat anti-rat (A488) or streptavidin A647 in blocking buffer at room temperature. All slides were then washed thrice in PBS before counterstained in Hoechst 33342 (1:1000) for 10 minutes. Goat anti-rabbit 488, goat anti-rabbit 633, goat anti-rat 488 were from Invitrogen; streptavidin 647 from Jackson ImmunoResearch. Slides were washed in PBS and then distilled water before mounted in mounting media for imaging by Zeiss LSM710 confocal microscope.

**Immunofluorescence quantifications**. For quantification of Cathepsin B intensity from immunostaining, four to six images per liver section were captured from different areas. The Cathepsin B mean fluorescence intensity (from Alexa fluor 488 signal) per image was measured using the same threshold by Fiji. Welch’s t-test was performed from 15 images for control and 18 images for gSpns1-cKO from three mice per genotype. For macrophages’ analysis, five to eight images per liver section were captured from different areas. The total number of cells per image was counted from Hoescht signal using Cell Profiler. Mac-2+ cells were indicated using Fiji, and the percentage of these cells was calculated per total cell number in each image. Around 3000 cells from each group were counted. Welch’s t-test was performed on 17 images for control and 20 images for gSpns1-cKO from three mice per genotype.

To quantify blood vessel density, embryonic brain sections (n=4 per genotype) were stained with GLUT1 antibody. The vascular density in the ganglionic eminence region was determined by Fiji. The immunofluorescence intensity of GLUT1 was quantified by Fiji. The diameters of cortex regions were also measured by Fiji (denoted by the line in Figure 1A).

**H&E staining**. Embryonic brain and adult liver sections from controls and gSpns1-cKO mice were prepared using a microtome and mounted onto microscope slides for overnight drying. The staining procedures have been described previously (9). Briefly, whole embryos collected at E13.5 and liver tissues collected from gSpns1-cKO and control mice were fixed in 4% PFA in phosphate buffer saline (PBS) at 4°C overnight. The next day, tissues were submerged in 15% sucrose for 24 hours and then 30% sucrose for 24 hours. The tissues were processed in a Leica tissue processor and then embedded in wax. Tissue sections were prepared using a microtome with 11-15 µm thickness. Serial sections were stained with hematoxylin and eosin. The slides were mounted in D.P.X mounting media and covered with coverslips. The sections were imaged using a Tissue scanner (FAXS) or Olympus microscope.

**Western blot**. To prepare the samples for WB analysis, embryos and tissues were homogenized in lysis buffer (with a ratio of 2-3 µl per 1 mg wet tissue) containing 150 mM NaCl, 1% Triton X-100, 10 mM Tris-HCl pH 8.0 with protease inhibitor (Thermofisher) and phosphatase inhibitor (Invitrogen) using FastPrep-24 MP with beads for 2 minutes in the cold room. Mammalian cells from HEK293 and CHO cell cultures were also lyzed in the lysis buffer for 30 minutes using a rotary shaker at 50 rpm in the cold room. Protein concentrations of lysates were measured using the BCA method. The same amounts of total proteins were added with Laemlli buffer and resolved in 10-12% SDS-PAGE at 100 V for 100 minutes. Proteins were transferred to Nitrocellulose membranes at 20 voltages in the cold room overnight. Membranes were then blocked in TBST (Tris-buffered saline with 0.1% Tween 20) containing 5% skim milk at room temperature for 1 hour. Primary antibodies diluted as advised by the manufacturer’s instruction in the same blocking buffer were incubated for 2 hours at room temperature or were incubated with the membranes in the cold room overnight. Next, the membranes were washed with TBST three times (5-minute intervals) before incubation in secondary antibodies (1:10,000) for 1 hour at room temperature. Membranes were then washed three times in TBST again before being visualized with ChemiDoc (Biorad).

**Enrichment of lysosomes from liver tissues.** Control and g*Spns1*-cKO mice were injected with 4 μl/g body weight of a 17% (w/v in 0.9% NaCl) Triton WR1339 solution (Sigma Aldrich) for three days prior to fractionation of lysosomes. The non-perfused livers were homogenized in five volumes (usually 5 ml) of ice-cold 0.25 M sucrose with three strokes in a 10 ml Potter-Elvejhem homogenizer. The homogenates were centrifuged at 1000 ×g for 10 minutes at 4°C. An amount of 4 ml supernatant was collected into 15 ml tubes, and the remaining pellet was resuspended in 3.5 ml 0.25 M sucrose, followed by another step of centrifugation at 1000 ×g for 10 minutes at 4°C for the second collection of 4 ml supernatant. Both supernatants (post-nuclear supernatant, PNS) were pooled and transferred to ultracentrifuge tubes. The supernatant was filled with 0.25 M sucrose up to 9 ml and centrifuged in an ultracentrifuge at 56,000 ×g (P40ST Swing rotor, Hitachi) for 7 minutes at 4°C. The supernatant was discarded. After resuspension of the pellet with 9 ml 0.25 M sucrose, followed by another step of ultracentrifugation at 56,000 ×g (P40ST Swing rotor, Hitachi) for 7 minutes at 4°C, the supernatant was also discarded. The pellet was resuspended in sucrose solution (3.5 ml) with a density of p=1.21 (resulting in the mitochondrial-lysosomal fraction (ML fraction). The ML fraction was transferred to an ultracentrifuge tube and overlaid by sequential steps with 2.25 ml of sucrose solutions of p=1.15, 2.25 ml of p=1.14, and 1 ml of p=1.06 yielding a discontinuous sucrose gradient. The discontinuous gradient was centrifuged at 110,000 ×g in a swinging bucket rotor (P40ST Swing rotor, Hitachi) for 150 minutes at 4°C. Lysosome fractions were collected at the interphase between p=1.14 and 1.06 sucrose (F2-fraction) after removing the top layer (F1 fraction).

**Internal standards for lipidomic analysis.** Internal standards (IS) solutions were prepared in butanol/methanol (BuMe) (1:1, v/v). The composition and concentration of the solutions were adjusted for the different sample types and analytical methods. For sphingolipid analysis, the internal standards Cer/Sph II from Avanti Polar Lipids (Cat LM 6005) were used. The IS in the sphingolipid extraction solvent contained 71.7 nM SM d18:1/12:0, 70.2 nM Sph d17:1, 71.4 nM Sph d17:0, 71.1 nM Cer and 69.9 nM d18:1/12:0 HexCer d18:1/12:0. For phospholipid analysis, the phospholipid IS for embryonic brain samples were used as follows: 1.18 µM PE (17:0/17:0), 2.5 µM DMPS, 3.12 µM PC 26:0 (13:0/13:0), 0.13 µM LPC 20:0, 0.165 µM LPE 14:0, 0.0525 µM DMPG, 0.34 µM PI 25:0, all of which were purchased from Avanti. The phospholipid IS for adult liver included 1.6 nM d3-Acyl Carnitine 16:0, 0.36 µM LPC 20:0, 5.57 µM PC 26:0 (13:0/13:0), 0.77 µM LPE 14:0, 3.01 µM PE (17:0/17:0), 0.75 µM DMPS, 0.83 µM PI 25:0, 0.15 µM DMPG, 0.90 µMSM 12:0 (d18:1/12:0). The phospholipid IS for lysosome fraction included SPLASH Mix (Avanti, 330707) (5.3 µM 15:0/18:1(d7) PC, 0.19 µM15:0/18:1(d7) PE, 0.13 µM 15:0/18:1(d7) PS (Na Salt), 0.93 µM 15:0/18:1(d7) PG (Na Salt), 0.27µM15:0/18:1(d7) PI (NH4 Salt), 0.27 µM 15:0/18:1(d7) PA (Na Salt), 1.20 µM 18:1(d7) Lyso PC, 0.27 µM18:1(d7) Lyso PE, 13.32 µM18:1(d7) Chol Ester, 0.13 µM18:1(d7) MAG, 0.40 µM 15:0/18:1(d7) DAG, 1.73 µM15:0/18:1(d7)-15:0 TAG, 1.07 µMd18:1/18:1(d9) SM, 6.66 µM Cholesterol (d7)).

**Methods for lipid extractions**

Embryonic brain and liver tissues: Embryonic brain tissues at E13.5 and PBS-perfused liver tissues from g*Spns1*-cKO and control mice were collected and immediately transferred to dry ice before storage at -80ºC. Tissue samples were then homogenized using FastPrep-24 MP with beads in 150 mM ammonium bicarbonate buffer, pH 7.8, with a ratio of 10 µl buffer per 1 mg wet tissue for 2 minutes in a cold room (4°C). The protein concentration of samples was measured using the BCA method. For lipid extraction, 200 µl of BuMe solution containing appropriate standards (phospholipid or sphingolipids, see above) were added to 20 µl of tissue homogenate. The IS-spiked samples were then sonicated in a water bath for 30 minutes, followed by a centrifugation at 14,000 ×g for 10 minutes. The supernatant was then transferred to mass spectrometry vials for LC-MS/MS analysis.

HEK293 and CHO cell: Cell samples were grown in 6-well plates, either in growth medium (DMEM with 10% fetal bovine serum, FBS) or starved under serum-deprived medium (DMEM without amino acids and FBS (Wako, 048-33575)) for 4 hours. Cells were harvested with cell scrappers using 1 ml PBS per well, then centrifuged at 3,000 ×g for 5 minutes to collect the cell pellets. An amount of 200 µl BuMe solution containing internal sphingolipid standard mix or phospholipid standard mix (see above) was added to the cell pellets. The IS spiked samples were then sonicated in a water bath for 30 minutes, followed by centrifugation at 14,000 ×g for 10 minutes. The supernatant was then transferred to mass spectrometry vials for LC-MS/MS analysis. The pellet was dried in Speedvac and then solubilized in 200 µl RIPA to measure protein concentration using BCA method for normalization.

Lysosome fractions: To extract sphingolipids from lysosomal fractions extracted from livers of control and gSpns1-cKO mice, 10 µl of each lysosomal fraction was used for lipid extraction using the above-mentioned procedure (addition of 200 µl BuMe containing the phospholipid internal standards). To extract phospholipids from lysosomal fraction, the knockout samples were first diluted 10 times in the same sucrose buffer (p=1.06), and then 20 µl of the lysosomal fraction from control and KO samples was used for lipid extraction as as described above (addition of 200 µl BuMe containing phospholipid internal standards, Splash Mix, Avanti). Protein concentrations in the lysosomal fractions were measured by the BCA method for normalization.

**Lipids quantification by LC/MS/MS**

Samples were randomized before extraction and analysis. Blank samples (empty tubes for cell samples and 10 µL MilliQ water for other sample types), matrix blanks (samples extracted without spiking internal standards), and pooled QC samples were used to assess method performance. In addition, diluted pooled QC samples were used to assess response linearity. Blanks, blank extracts, QC, and diluted QC were interspersed with study samples throughout the analytical run.

Sphingolipids were separated using a reverse phase column (Agilent RRHD Eclipse Plus column, 959758-902, C18, 2.1x100 mm, 1.8 µm) on an Agilent 6495A2 Triple Quadrupole mass spectrometer (Agilent Technologies). Mobile phases A (60% Methanol (Thermo Fisher Scientific Inc.), 40% MilliQ water, 10 mM Ammonium Acetate (Sigma-Aldrich), 0.2% formic acid (Sigma-Aldrich)) and B (60% Methanol (Thermo Fisher Scientific Inc.), 40% Isopropanol (Thermo Fisher Scientific Inc.), 10 mM Ammonium Acetate (Sigma-Aldrich), 0.2% formic acid (Sigma-Aldrich)) were mixed using the following gradient: 0–3 minutes, 0-10% B; 3-5 minutes, 10-40% B; 5-5.3 minutes, 40-55% B; 5.3-8 minutes, 55-60% B, 8-8.5 minutes, 60-80% B; 8.5-10.5 minutes, 80% B; 10.5-16 minutes, 80-90% B; 16-19 minutes, 90% B, 19-22 minutes, 90-100% B. The flow rate was 0.4 ml/min, and the sample injection volume was 2 µl.

Phospholipids were separated using a HILIC column (Kinetex 2.6 µm HILIC 100 Å, 150 × 2.1 mm, Phenomenex, Torrance, CA, USA) on an Agilent 6495A2 (Agilent Technologies). Mobile phases A (50% acetonitrile (LC-MS grade, Thermo Fisher Scientific Inc.) 50% 25 mM ammonium formate (Sigma-Aldrich) pH = 4.6) and B (95% acetonitrile and 5% 25 mM ammonium formate pH = 4.6) were mixed at the following gradient: 0–6 minutes, 99–75% B; 6–7 minutes, 75-10% B; 7–7.1 minutes, 10–99.9% B; 7.1–10.1 minutes, 99.9% B. The flow rate was 0.5 ml/minute, and the sample injection volume was 0.5 µL for lysosome fraction and 0.25 µL for liver samples.

The MS parameters on Agilent 6495A2 were as follows: electrospray ionization, gas temperature 200°C, gas flow 15 l/minute, sheath gas flow 12 l/minute, and capillary voltage 3,500 V. Sphingolipids and phospholipids were quantified at the sum composition level using multiple reaction monitoring (MRM), see supplementary table (Supplementary table S15) for detailed MRMs.

**Data Processing for lipidomic**. The raw data were processed using Agilent MassHunter Quantitative Analysis software for QQQ version B.08 (Agilent Technologies, Santa Clara, CA, USA). The areas under the curve (AUC) were manually inspected and checked for retention time. Blanks and QC samples were used to assess the following parameters for each MRM transition: coefficient of variation (CV) in QCs, signal in blanks, and linearity in diluted QC samples. Only MRM transitions satisfying the established criteria were used for quantification: CV in QCs < 25%, signal QC/blank > 10, and Pearson R2 in diluted QC > 0.8. Isotopic correction of signals based on the precursor ions was performed using LICAR (Gao et al., Anal. Chem. 2021,93,3163) for HILIC analysis. Finally, the lipid samples were normalized to relevant internal standards and protein concentration.

**Metabolomics**

For metabolomics, we collected 50-100 mg tissues from PBS perfused livers of *Spns1^f/f^*-Rosa26CreERT^2^ (g*Spns1*-cKO) and control mice. The untargeted metabolomics was performed by Metabolon. Detail methods for metabolomic analysis from Metabolon is described below.

For sample preparation: Liver samples were prepared using the automated MicroLab STAR® system from Hamilton Company. Several recovery standards were added prior to the first step in the extraction process for QC purposes. To remove protein, dissociate small molecules bound to protein or trapped in the precipitated protein matrix, and to recover chemically diverse metabolites, proteins were precipitated with methanol under vigorous shaking for 2 min (Glen Mills GenoGrinder 2000) followed by centrifugation. The resulting extract was divided into five fractions: two for analysis by two separate reverse phases (RP)/UPLC-MS/MS methods with positive ion mode electrospray ionization (ESI), one for analysis by RP/UPLC-MS/MS with negative ion mode ESI, one for analysis by HILIC/UPLC-MS/MS with negative ion mode ESI, and one sample was reserved for backup. Samples were placed briefly on a TurboVap® (Zymark) to remove the organic solvent. The sample extracts were stored overnight under nitrogen before preparation for analysis. For metabolite analysis, ultrahigh Performance Liquid Chromatography-Tandem Mass Spectroscopy (UPLC-MS/MS) was employed. All methods utilized a Waters ACQUITY ultra-performance liquid chromatography (UPLC) and a Thermo Scientific Q-Exactive high resolution/accurate mass spectrometer interfaced with a heated electrospray ionization (HESI-II) source and Orbitrap mass analyzer operated at 35,000 mass resolution. Prior to analysis, the sample extract was dried then reconstituted in solvents compatible to each of the four methods. Each reconstitution solvent contained a series of standards at fixed concentrations to ensure injection and chromatographic consistency. One aliquot was analyzed using acidic positive ion conditions, chromatographically optimized for more hydrophilic compounds. In this method, the extract was gradient eluted from a C18 column (Waters UPLC BEH C18-2.1x100 mm, 1.7 µm) using water and methanol, containing 0.05% perfluoropentanoic acid (PFPA) and 0.1% formic acid (FA). Another aliquot was also analyzed using acidic positive ion conditions; however, it was chromatographically optimized for more hydrophobic compounds. In this method, the extract was gradient eluted from the same afore mentioned C18 column using methanol, acetonitrile, water, 0.05% PFPA and 0.01% FA and was operated at an overall higher organic content. Another aliquot was analyzed using basic negative ion optimized conditions using a separate dedicated C18 column. The basic extracts were gradient eluted from the column using methanol and water, however with 6.5mM Ammonium Bicarbonate at pH 8. The fourth aliquot was analyzed via negative ionization following elution from a HILIC column (Waters UPLC BEH Amide 2.1 x 150 mm, 1.7 µm) using a gradient consisting of water and acetonitrile with 10mM Ammonium Formate, pH 10.8. The MS analysis alternated between MS and data-dependent MS^n^ scans using dynamic exclusion. The scan range varied slighted between methods but covered 70-1000 m/z. Raw data files are archived and extracted as described below.

**Data Extraction and Compound Identification from metabolomics.** Raw data was extracted, peak-identified and QC processed using Metabolon’s hardware and software. These systems are built on a web-service platform utilizing Microsoft’s .NET technologies, which run on high-performance application servers and fiber-channel storage arrays in clusters to provide active failover and load-balancing. Compounds were identified by comparison to library entries of purified standards or recurrent unknown entities. Metabolon maintains a library based on authenticated standards that contains the retention time/index (RI), mass to charge ratio (*m/z)*, and chromatographic data (including MS/MS spectral data) on all molecules present in the library. Furthermore, biochemical identifications are based on three criteria: retention index within a narrow RI window of the proposed identification, accurate mass match to the library +/- 10 ppm, and the MS/MS forward and reverse scores between the experimental data and authentic standards. The MS/MS scores are based on a comparison of the ions present in the experimental spectrum to the ions present in the library spectrum. While there may be similarities between these molecules based on one of these factors, the use of all three data points can be utilized to distinguish and differentiate biochemicals. More than 3300 commercially available purified standard compounds have been acquired and registered into LIMS for analysis on all platforms for determination of their analytical characteristics. Additional mass spectral entries have been created for structurally unnamed biochemicals, which have been identified by virtue of their recurrent nature (both chromatographic and mass spectral). These compounds have the potential to be identified by future acquisition of a matching purified standard or by classical structural analysis.

**Metabolite curation**. A variety of curation procedures were carried out to ensure that a high-quality data set was made available for statistical analysis and data interpretation. The QC and curation processes were designed to ensure accurate and consistent identification of true chemical entities, and to remove those representing system artifacts, mis-assignments, and background noise. Metabolon data analysts use proprietary visualization and interpretation software to confirm the consistency of peak identification among the various samples. Library matches for each compound were checked for each sample and corrected if necessary.

**Metabolite Quantification and Data Normalization.** Peaks were quantified using area-under-the-curve. For studies spanning multiple days, a data normalization step was performed to correct variation resulting from instrument inter-day tuning differences. Essentially, each compound was corrected in run-day blocks by registering the medians to equal one (1.00) and normalizing each data point proportionately (termed the “block correction”; Figure 2). For studies that did not require more than one day of analysis, no normalization is necessary, other than for purposes of data visualization. In certain instances, biochemical data may have been normalized to an additional factor (e.g., cell counts, total protein as determined by Bradford assay, osmolality, etc.) to account for differences in metabolite levels due to differences in the amount of material present in each sample.

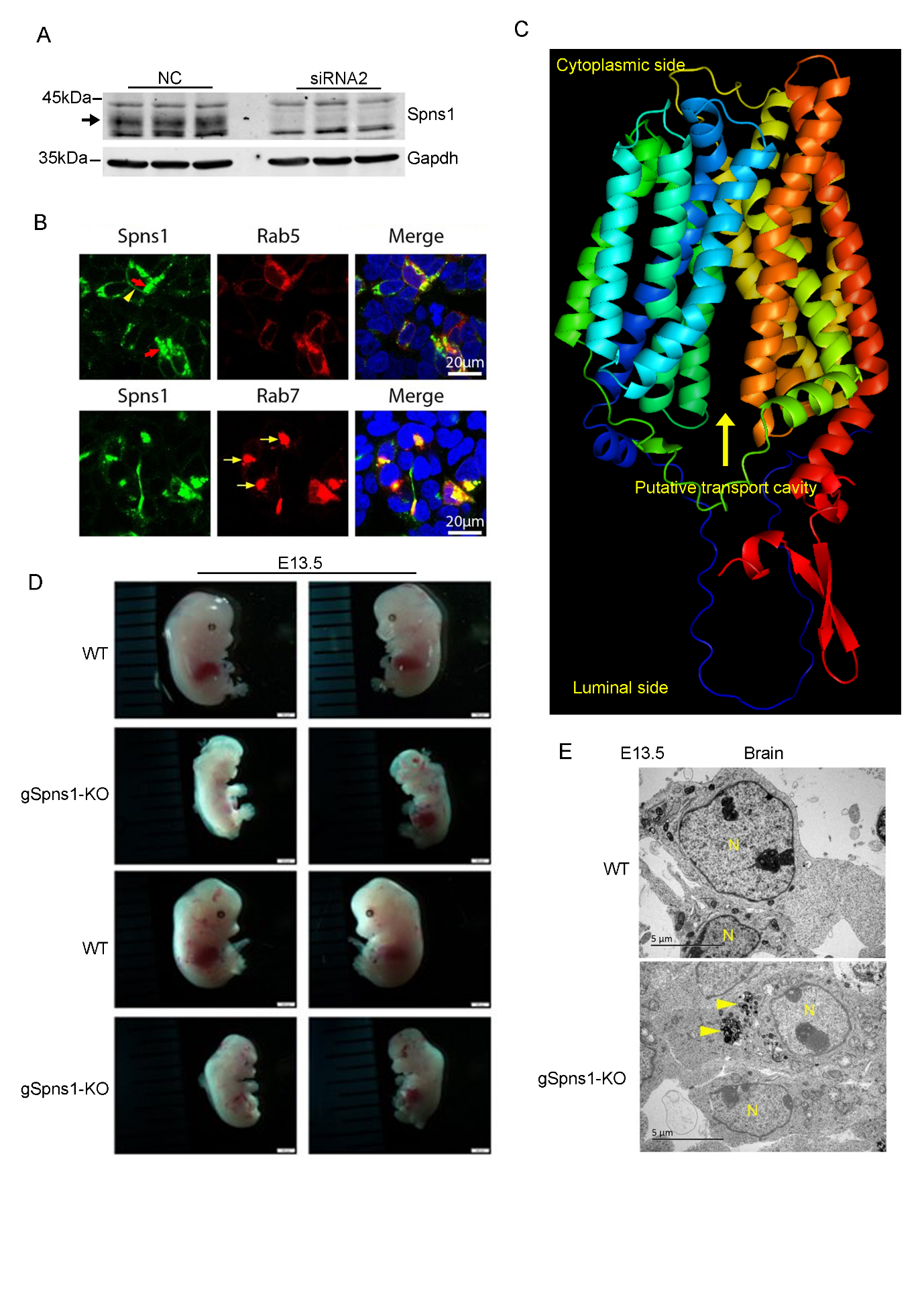

***Figure S1. SPNS1 is a putative lysosomal transporter required for development****.* ***A****, Validation of polyclonal antibodies for SPNS1 in CHO cells. Expression of SPNS1 in wild-type and Spns1 knockdown CHO cells. The data show that endogenous SPNS1 is expressed in CHO cells with a molecular weight of approximately 40 kDa.* ***B****, SPNS1 is co-localized with late endosomal maker Rab7 (lower panel, arrows), but not with early endosomal marker Rab5 (upper panel, arrows). SPNS1 is co-expressed with Rab7-RPF or Rab5-RFP in HEK293 cells. SPNS1 is present in or near plasma membrane (upper panel, arrowhead).* ***C****, Modelled structure of human Spns1 from AlphaFold2. Arrow shows putative transport cavity predicted for a solute transporter.* ***D****, Whole body deletion of Spns1 results in early lethality in mice. Shown are representative images from 2 WT and 2 KO embryos of the same litters. The whole body of Spns1 knockout is smaller with severe brain development.* ***E****, Electron microscopic analysis of brain sections from E13.5 WT and Spns1 knockout embryos. Representative images from WT and KO embryos.* *Knockout embryos had accumulation of condensed membranous materials in the cytoplasm (arrowheads). N, nuclei. n=3 per genotype.*

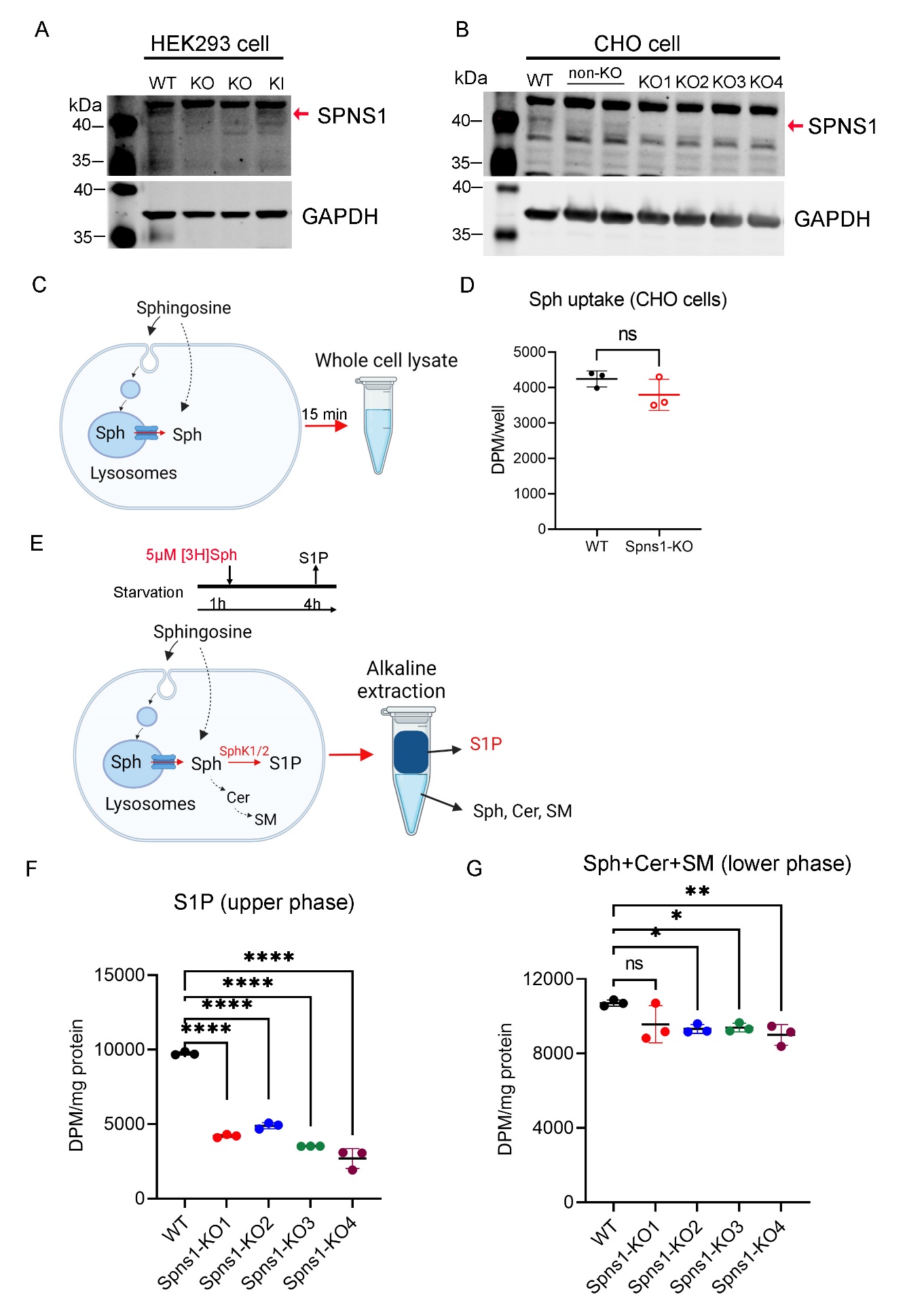

***Figure S2. SPNS1 is required for sphingosine release from lysosomes. A-B,*** *Generation of Spns1 knockout and knock-in cells by CRISPR/Cas9 technology. Western blot analysis of SPNS1 protein band in wild-type, Spns1 knockout, and Spns1 knock-in in HEK293 cells (in* ***A****) and in Spns1-KO clones in CHO cells (in* ***B****).* ***C-D****, Whole-cell sphingosine uptake assay in WT and Spns1-KO CHO cells. Total radioactive signals were measured to quantify the levels of sphingosine import* *in WT and Spns1-KO CHO cells after incubation with [3-^3^H]-sphingosine for 15 minutes.* ***E****, Illustration of [3-^3^H]-sphingosine transport assays. CHO cells were starved in medium without amino acids and serum for 1 hour. The cells were then added with radioactive sphingosine and continued incubating for 4 hours.* ***F****. Radioactive S1P levels (upper phase) from WT and different Spns1-KO clones from CHO cells.* ***G****, Radioactive sphingolipid levels (lower phase) from WT and different Spns1-KO clones. *p<0.05; **p<0.01; ****p<0.0001; ns, not significant. Data are expressed as mean ± SD. Statistical significance was determined by* o*ne-way ANOVA.*

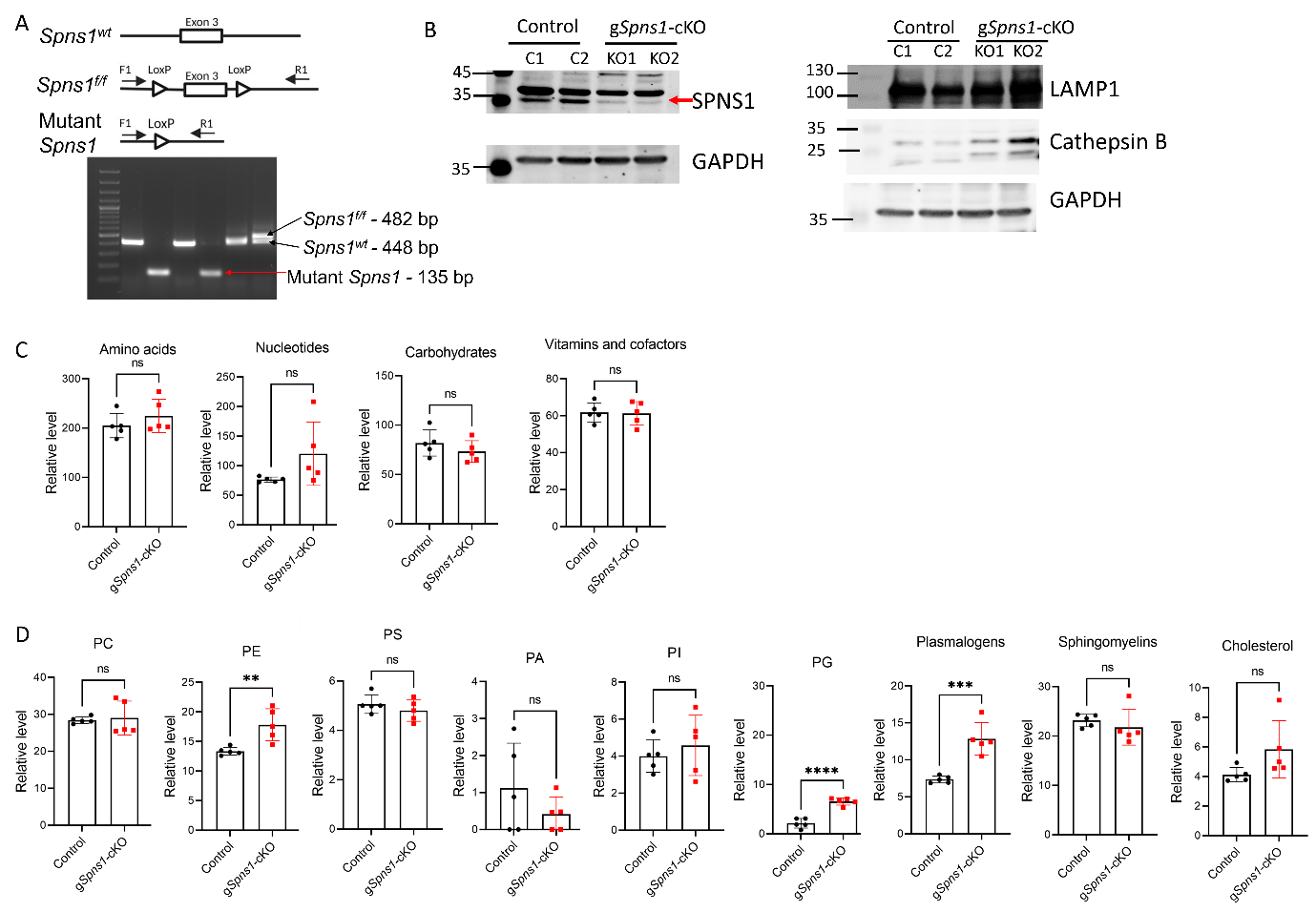

***Figure S3. Postnatal deletion of Spns1 does not affect the levels of major phospholipids such as glycerophospholipids and sphingolipids in the liver. A,*** *Generation of conditional knockout of Spns1 by CRISPR/Cas9 technology****. B,*** *Validation of Spns1 deletion in livers of control and gSpns1-cKO mice (Spns1f/f; ROSA26 CreER^T2^ mice) after induction with tamoxifen. Expression of SPNS1 in post nuclear fractions (PNS) from livers of control and gSpns1-cKO mice. Arrow shows the reduction of Spns1 protein band in the knockouts. n=2 per genotype.* ***C,*** *Metabolomic analysis of amino acids, nucleotides, carbohydrates, and vitamins and cofactors from livers of control and gSpns1-cKO mice.* ***D****, Metabolomic analysis of PC, PE, PA, PI, plasmalogens, PS, sphingomyelins, and cholesterols in the livers of control and gSpns1-cKO mice. **p<0.01;* ****p<0.001; ****p<0.0001; ns, not significant. Data are expressed as mean ± SD. Statistical significance was determined by two-sided unpaired t-test.*

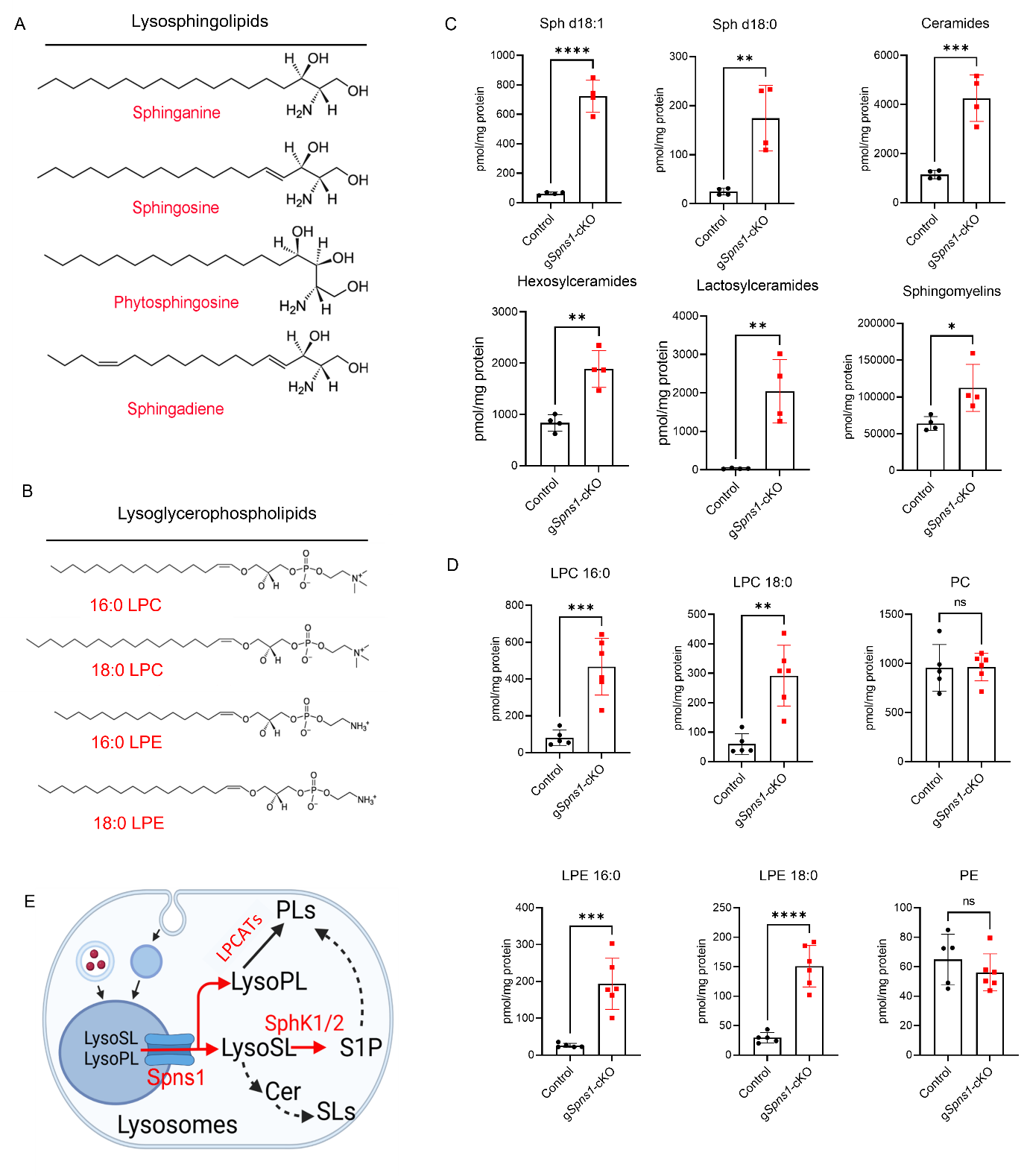
***Figure S4. Deletion of Spns1 results in accumulation of major lysophospholipids. A-B,*** *Chemical structure of major lysosphingolipids (A) and lysoglycerophospholipids (B).* ***C,*** *Total levels of major sphingolipids in the livers of control and gSpns1-cKO mice by targeted lipidomics. Each symbol represents one mouse.* ***D****, Total levels of major phospholipids such as PC, PE in the livers of control and gSpns1-cKO mice by targeted lipidomics. Each symbol represents one mouse.* ***E****, Proposed model of lysosomal transport of lysoglycerophospholipids including lysosphingolipids (lysoSL) and lysoglycerophospholipids (lysoPL) by SPNS1. PLs, phospholipids; SLs, sphingolipids; Cer, ceramides. *p<0.05, **p<0.01, ***p<0.001, ****p<0.0001; ns, not significant, Data are expressed as mean ± SD. Statistical significance was determined by two-sided unpaired t-test.*

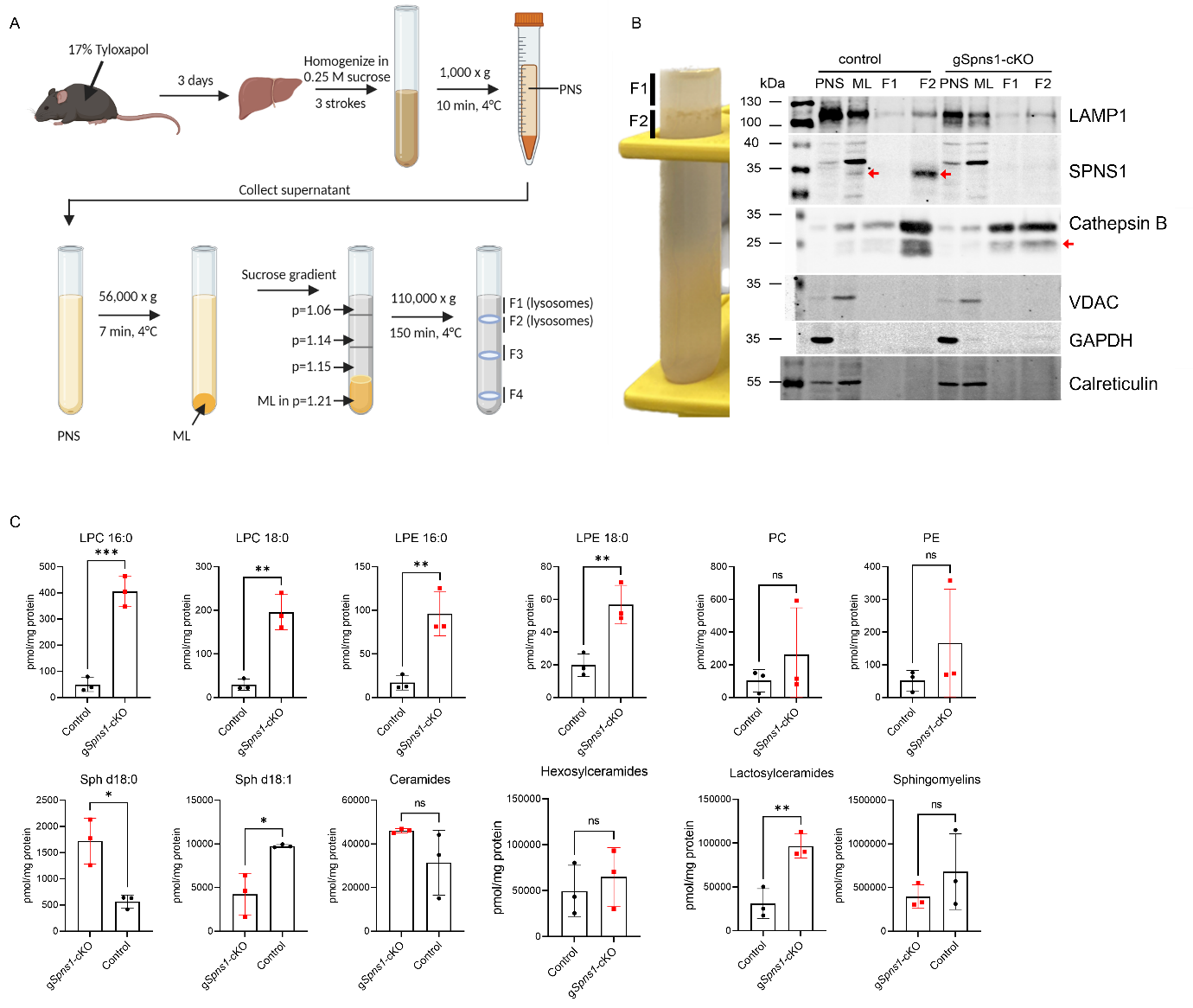
***Figure S5. Lack of Spns1 brings about phenotypes reminiscent of sphingolipid lysosomal storage diseases****.* ***A****, Schematic illustration of isolation of lysosomes from mouse liver tissues.* ***B****, Western blot analysis of lysosomal (LAMP1 and Cathepsin B), mitochondria (Vdac), and endoplasmic reticulum (Calreticulin) markers from liver lysosome fractions.* ***C****, The levels of sphingolipids and phospholipids from lysosome fractions from livers of control and gSpns1-cKO mice. n=3 per genotype. *p<0.05; **p<0.01; ***p<0.001; ns, not significant, Data are expressed as mean ± SD. Statistical significance was determined by two-sided unpaired t-test.*

**Legends for supplemental tables.**

**Supplemental table S1:** Lipidomic analysis of sphingolipids from brains of WT and g*Spns1*-KO embryos.

**Supplemental table S2:** Lipidomic analysis of sphingolipids from WT and SPNS1-KO in HEK293 cells.

**Supplemental table S3:** Lipidomic analysis of sphingolipids from WT and SPNS1-KO in CHO cells.

**Supplemental table S4:** Lipidomic analysis of sphingolipids from WT, SPNS1-KO, and NPC1 knockout in HEK cells.

**Supplemental table S5:** Data used for 'Figure 3A - Liver: Lysophospholipids' and 'Figure 3H - Liver: Sphingolipids'.

**Supplemental table S6:** Total levels of different phospholipids and cholesterol metabolites.

**Supplemental table S7:** Metabolomic analysis results for all lipid metabolites.

**Supplemental table S8:** Levels of amino acid metabolites.

**Supplemental table S9:** Levels of carbohydrate metabolites.

**Supplemental table S10:** Levels of nucleotide, vitamin, and cofactor metabolites.

**Supplemental table S11:** Sphingolipid (SL) analysis of whole liver tissues from control and g*Spns1*-cKO mice.

**Supplemental table S12:** Phospholipid (PL) analysis of whole liver tissues in g*Spns1*-cKO mice.

**Supplemental table S13:** Sphingolipid analysis of lysosome extraction (combined F1 and F2 fractions) from liver tissues of control and gSpns1-cKO mice.

**Supplemental table S14:** Phospholipid analysis of lysosome extraction (combined F1 and F2 fractions) from liver tissues of control and gSpns1-cKO mice.
